# SPARC regulation of PMN clearance protects from pristane induced lupus and rheumatoid arthritis

**DOI:** 10.1101/2020.08.10.243816

**Authors:** Sabina Sangaletti, Laura Botti, Alessandro Gulino, Daniele Lecis, Barbara Bassani, Paola Portararo, Matteo Milani, Loris De Cecco, Matteo Dugo, Claudio Tripodo, Mario P. Colombo

## Abstract

One step along the pathogenesis of Systemic lupus erythematosus (SLE) is associated with polymorphonuclear leukocyte (PMN) death and their ineffective removal by M2-macrophages. The secreted protein acidic and rich in cysteine (SPARC) is a matricellular protein with unexpected immunosuppressive function in M2-macrophages and myeloid cells. To investigate the role of SPARC in autoimmunity, we adopted a pristane–induced model of lupus in mice, which recapitulates clinical manifestations of human SLE. *Sparc*^*-/-*^ mice developed earlier and more severe renal disease, lung and liver parenchymal damage than the WT counterpart. Most prominently, *Sparc*^*-/-*^ mice had anticipated and severe occurrence of arthritis. An intermediate phenotype was obtained in *Sparc*^*+/-*^ hemizygous mice, a result that suggests *Sparc* gene-dosage as relevant in autoimmune-related events. Mechanistically, a defective *Sparc* expression in PMN blocks their clearance by macrophages, through a defective delivery of eat-me and don’t eat-me signals. *Sparc-/-* PMN that escape macrophage scavenging becomes a source of autoantigens for dendritic cell (DC) presentation and a direct stimulus for IL-17 expression in γδ-T-cells. Gene profile analysis of synovial biopsies of knees affected by SLE-associated arthritis showed an inverse correlation between SPARC and key autoimmune genes. These results point to SPARC down-regulation as a key event characterizing SLE and associated rheumatoid arthritis pathogenesis.

## Introduction

Autoimmunity results from multiple events dysregulating immune responses toward abnormal recognition of self-antigens, loss of tolerance, and the consequent attack of healthy cells and tissues.

The pathogenesis of autoimmune diseases combines genetic and epigenetic factors with environmental stressors, such as persistent inflammatory conditions, infections, and tissue damage ((Costenbader and Karlson, 2006; Rosenblum et al., 2015)).

Systemic lupus erythematosus (SLE) is a prototypical highly complex autoimmune disease characterized by circulating autoantibodies to nuclear and cytoplasmic antigens and by widespread involvement of the skin, joints, and parenchymal organs (Dorner and Furie, 2019). Although autoantibodies to DNA are a consistent finding in SLE, their direct role in determining the tissue/organ damage has been revised in favor of a more complex interaction between innate and adaptive autoimmunity (Bagavant and Fu, 2005). In systemic vasculitis (SV) clinical and experimental evidences support a pathogenesis driven by anti-neutrophil cytoplasmic autoantibodies (ANCAs). In SV, ANCAs induce neutrophil extracellular traps (NETs) (Heeringa et al., 2018) that, composed by extracellular DNA threads decorated with proteolytic and potentially injuring enzymes, directly promote endothelial damage (Kessenbrock et al., 2009). Moreover, the uptake by dendritic cells (DC) of NET-coated with antigens recognized by ANCA-Abs, such as myeloperoxidase (MPO) and proteinase-3, begin and perpetuate the autoimmune responses by promoting ANCA induction (Sangaletti et al., 2012). In SLE, NETs are potent stimulators of pathogenic plasmacytoid dendritic cells (pDCs) ((Garcia-Romo et al., 2011; Lande et al., 2011) and direct activators of human autoreactive memory B cells (Gestermann et al., 2018). Therefore, any unbalance in NET formation can be linked with the pathogenesis of SLE and SV. In SLE, neutrophils are characterized not only by exacerbated NETosis, but also by defective phagocytosis, microbial killing capacity and cell debris removal, by increased oxidative stress and cytokine production, and enhanced apoptosis (Tsai et al., 2019) (Kaplan, 2011). These defects are magnified by other phagocytic cell impairment, including defective macrophages efferocytosis. Indeed, patients with SLE often display a deficiency in clearing apoptotic cells and immune complexes (ICs) ((Munoz et al., 2010) (Baumann et al., 2002) (Davies et al., 2002)) and also defective expression of monocytes adhesion molecules like CD44 (Cairns et al., 2001).

The secreted protein acidic and rich in cysteine (SPARC) is a multi-faceted matricellular protein involved in normal and pathological tissue remodeling (Puolakkainen et al., 2003; Trombetta-Esilva and Bradshaw, 2012) and capable of affecting, either positively or negatively, cell-cell and cell-ECM adhesive interactions (Brekken and Sage, 2001). Some divergent functions of SPARC reveal its complexity and underscore the context dependency of its activity, which strictly depends on tissue and cellular origin (Bradshaw, 2012). A under-investigated but interesting aspect of SPARC function, is the capacity of regulating the immune system in several aspects, such as dendritic cell migration and branching (Piconese et al., 2011), myeloid-derived suppressor cell expansion and functional differentiation (Sangaletti et al., 2019) and B cell development (Luo et al., 2014; Sangaletti et al., 2014a). Studying mesenchymal cells functions in bone-marrow (BM) and secondary lymphoid organs we showed that the defective B cells development occurring in *Sparc*^*-/-*^ hosts under radiation-induced stress relies on SPARC limiting B cells precursors apoptosis (Sangaletti et al., 2014a). Furthermore, in Fas mutant mice we showed that neutrophils from *Sparc*^*-/-*^ mice were more supportive for B-cell transformation through an increased expression of BAFF and IL-21 and increased NETosis(Sangaletti et al., 2014b). These evidences suggested an involvement of SPARC in the regulation of immune cell death, a key immunological checkpoint to prevent autoimmunity.

To investigate the functional role of SPARC in SLE, we adopted a chemically– induced model of autoimmunity, namely the intraperitoneal (i.p.) injection of the hydrocarbon oil 2,6,10,14-tetramethylpentadecane (TMPD), more commonly known as pristane, which induces a systemic lupus-like disease, in BALB/c mice. This model recapitulates several major clinical manifestations of human lupus including glomerulonephritis, lung vasculitis and arthritis, accumulation of autoantibodies to single (ss)- and double (ds)-strand-DNA, to chromatin and ribonucleproteins (RNP), as well as the female-sex biased incidence of the disease and the dependency on IFN for disease initiation (Reeves et al., 2009).

## Material and Methods

### Mice and treatment

*Sparc*^*-/-*^ and WT mice were on BALB/c background and were bred and maintained in our animal facility. In some experiments WT mice were purchased from Charles River. Experiments were performed according to local ethical guidelines. Authorization number (internal ethical committee): INT_10/2011).

8-10 weeks animals were intraperitoneally injected with 0,5 ml of Pristane (2,6,10,14-Tetramethylpentadecane, Sigma-Aldrich) at day 0, 60 and 120 and monitored every two weeks for signs of arthritis. Severity of arthritis was evaluated in a blind fashion for each paw per mouse with a 0-3 score (maximum 12 pt each animal): 0, no arthritis; 1, mild joint swelling and erythema; 2, severe joint and digit swelling; 3, paw deformation and ankylosis.

For Collagen Antibody Induced Arthritis (CAIA), 2 mg of ArthritoMab Antibody cocktail were administered i.v. on day 0 and 50 μg LPS was administered i.p. on day 3 (All from MDBiosciences). All four paws were scored daily using a 0-3 score.

### Cell preparations

Peritoneal lavage was performed by infusing 10 ml of sterile PBS. Blood was collected in heparinized tubes, erythrocytes were lysed. Two popliteal lymph nodes from each animal were pooled and mechanically disrupted in PBS; suspensions were passed through a 70 μm cell strainer and used for subsequent analysis. Cells from paws were obtained as previously described (Pollinger et al., 2011). Briefly, hind paws from each mouse were cut, skin was removed and the paws were chopped into small pieces. Pieces were resuspended in HBSS and digested in 1mg/ml collagenase IV (Sigma-Aldrich) at 37 °C for 1 hour; digested tissue were passed through 70 μm cell strainer and cell suspension were used for subsequent analysis.

### Flow cytometry

Surface staining was performed on single cell suspension using fluorochrome-conjugated monoclonal antibodies against CD4, B220 or CD19, CD11b, Gr1/Ly6G/C, F4/80, CD31, CD47, CD44, CD62L, CD69, CD11c (eBioscience), calreticulin (Abcam). AnnexinV /7AAD staining was performed using products and instructions from eBioscience. For intracellular staining, cells were first stimulated 4 hours *in vitro* with Cell Stimulation Cocktail (plus protein transport inhibitors) (eBioscience). Then cells were surface stained as above and fixed 20 minutes using the IC fixation buffer, followed by incubation with monoclonal antibodies against IL-17, TNF, IFN-γ and resuspendend in Permeabilization Buffer (all from eBioscience). Samples were acquired on the LSR II (BD Biosciences) and analyses were performed using FlowJo software (TreeStar).

### PMN Activation and ROS measurement

Peritoneal cavity (PerC) cells of WT and *Sparc*^*-/-*^ were seeded in 96 well plates and incubated 5 minutes at 37 °C with Dihydrorhodamine 123 (DHR 123) 5 μg/ml in medium, washed and incubated with APC conjugated anti mouse Gr1 for 15’. After washing, cells were activated using phorbol-12-myristate-13-acetate (PMA) 100 ng/ml for the indicated time, after which activation was blocked immediately by diluting samples with fresh medium and keeping cells in ice. Samples were acquired immediately using BD LSR Fortessa; PMN were identified using physical parameters and Gr1 expression and mean fluorescence intensity (MFI) of Rhodamine123 positive cells was calculated.

### In vivo phagocytosis assay

To evaluate phagocytosis activity of PerC macrophages, WT or *Sparc*^*-/-*^ mice were treated with 1 ml thioglycollate i.p. 4 days later, CFSE labeled Jurkat cells were UV irradiated for 20 minutes to induce apoptosis and injected in the PerC of treated mice. After 30 minutes, PerC were washed and the frequency of Mϕ that have engulfed CFSE+ apoptotic bodies were examined and indicated as % of phagocytizing F4/80^high^ and F4/80 ^int^ Mϕ.

### Gene expression profiling analysis

PerC macrophages obtained from WT or Sparc-/- mice were extracted using RNAeasy micro kit (Qiagen). After quality check and quantification by 2100 Bioanalyzer system (Agilent) and Nanodrop ND-1000 spectrophotometer (ThermoFisher), respectively, gene expression was assessed using Illumina BeadChip following manufacture’s protocol.

Raw expression data were collected from scanned images using Illumina BeadStudio v3.3.8 and processed using the *lumi* package from Bioconductor. Raw data were log_2_-transformed, normalized with robust spline normalization and filtered, keeping only probes with a detection *p*-value < 0.01 in at least one sample. Multiple probes mapping to the same gene symbol were collapsed using the collapse Rows function of the WCGNA package with “MaxMean” method. Expression data were deposited in NCBI’s Gene Expression Omnibus (GEO) database with accession number GSE144415.

Differentially expressed genes were identified using *limma* package. *P*-values were corrected for multiple testing using the Benjamini-Hochberg false discovery rate (FDR) method. An FDR < 0.25 was applied to select differentially expressed genes. Gene Set Enrichment Analysis (GSEA) was performed using the fgsea package and hallmark gene sets collected from the MSigDB database (http://www.broadinstitute.org/gsea/msigdb/index.jsp) by ranking genes according to their t-statistic. Significantly enriched gene sets were selected according to an FDR < 0.05.

We selected GSE36700 dataset from NCBI, which contained gene expression of synovial biopsies from a total of 25 patients affected by: Rheumatoid Arthritis (RA), Seronegative Arthritis (SA), Systemic Lupus Erythematosus (SLE) and Osteoarthritis (OA). Those samples were profiled with Affymetrix Human Genome U133 Plus 2.0 microarrays. Normalized data were collected from GEO repository, probes without their corresponding gene symbol were removed and multiple probes associated to the same gene symbol were collapsed using the collapseRows function of the WCGNA package with “maxRowVariance” method. To discover an overall change of expression in each class of disease a selected list of genes was tested with ANOVA and the variation was considered significant with P-Value < 0.05. Significant pair-wise differences (P-Value < 0.05) were identified using the post-hoc Tukey’s honest significance test. Single sample gene set enrichment analysis (ssGSEA) was performed with the GSVA package for two hallmark pathways retrieved from MSigDB database: epithelial mesenchymal transition and interferon gamma response; then t-test was computed to find intra-disease gene set variations and comparison was considered significant with the false discovery rate (FDR) < 0.05, calculated with the Benjamini & Hochberg correction.

## Results

### SPARC protects mice from onset of SLE-like disease

Mice treated with pristane develop a SLE-like disease characterized by multiorgan damage and auto-antibodies (Reeves et al., 2009). We injected pristane i.p. in wild-type (WT) and *Sparc*^*-/-*^ mice that were monitored over a 300 days-period. Histopathological analysis of the kidney revealed that differently from WT mice, which displayed glomerular atrophy, *Sparc*^*-/-*^ mice developed earlier and more severe renal disease with glomerular hypercellularity, sclerosis with the presence of immune infiltrates, lipoid tubular degeneration and also glomerular necrosis (Figure 1A, right, inset). *Sparc*^*-/-*^ mice also showed more severe lung injury on histopathology, with signs of alveolar hemorrhage and massive hemosiderin-laden histiocytic infiltration (Figure 1B). In addition to kidney and lung damage, the liver of *Sparc*^*-/-*^ mice showed more severe signs of portitis evolving to steatohepatitis than WT liver counterpart (Figure 1C). However, the most prominent phenotype due to SPARC deficiency was detected in the joints. Indeed, *Sparc*^*-/-*^ mice had increased and anticipated occurrence of arthritis (130 days vs 220 days) (Figure 1D-E) in one or more paws and more severe manifestations of arthritis than the *Sparc* proficient counterpart (Figure 1F). Histological examination of arthritic damage at 130 and 220 days post pristane injection showed that paws from *Sparc*^*-/-*^ mice were heavily infiltrated by a mixed population of immune cells mostly composed by polymorphonuclear granulocytes and, to a lesser extent, lymphocytes (Figure 1G).

**Figure 1:**
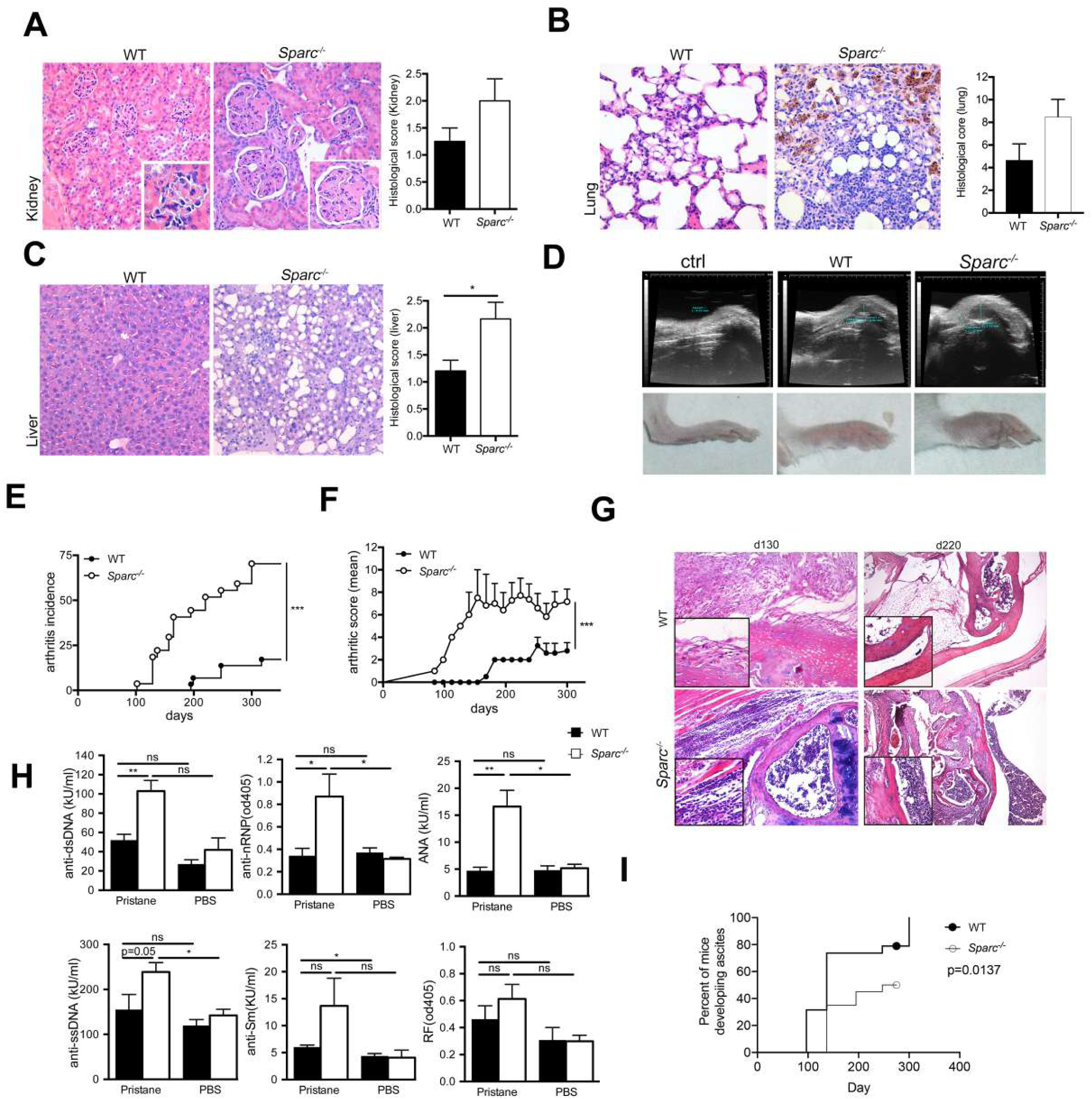
SPARC deficiency causes accelerated production of autoantibody and exacerbated organ damage. A) Representative histology of kidneys (20X) from WT and *Sparc*^*-/-*^ mice 300 days after initiation of the experiment showing more severe organ damage of the latter group of animals (left) and histological score evaluated on 5 mice per group (right); inset, a glomerulus at higher magnification (40X); B) representative histology (left) and histological score (right) of lung from the indicated mice (20X); C) As in A, representative image and score of liver (20X); D) The upper panel shows a representative ultrasound (US) echographic examination of the arthritic paw of pristane-treated WT and *Sparc*^*-/-*^ mice. The lower panel shows representative pictures of paws from WT and *Sparc*^*-/-*^ at day 200 after first pristane injection; E) Kinetic analysis of arthritis incidence; F) Arthritis score obtained evaluating pristane treated mice every two weeks (n= 17 mice/group pooled from two independent experiments); G) Representative H&E staining of arthritic paws at day 130 and 220 after pristane injection; H) Total serum Igs against the indicated self antigens have been dosed in untreated mice or mice 130 days after pristane treatment (n=7 for treated mice and n=4 for untreated controls). One experiment representative of three is shown. I. Ascites development in WT and *Sparc*^*-/-*^ mice.

Consistent with the more severe phenotype occurring in *Sparc*^*-/-*^ mice, their sera, evaluated 130 days post pristane, were characterized by the enrichment in autoantibodies against self-nuclear antigens (Figure 1H), including dsDNA, nRNP, ANA. Notably, at the same time point, the *Sparc* proficient WT counterpart did not show any significant production of autoantibodies. The difference was equalized at later time points being the level of autoantibodies very similar 200 days post pristane treatment (not shown). This suggests an anticipated onset of auto-antibody production in *Sparc-/-* mice. Of note, the onset of this phenotype was independent from the development of ascites that was reduced in *Sparc-/-* rather than in WT mice (Figure 1I).

### Pristane-elicited PMN undergo accelerated cell death in Sparc^-/-^ mice

We reasoned that one key event that might explain the earlier autoantibody production occurring in *Sparc-/-* mice could be ascribed to an increased and early availability of auto-antigens. The injection of pristane in mice leads to a persistent inflammatory response in the peritoneal cavity, with massive recruitment of polymorphonuclear leukocytes (PMN) that represent, upon death, a potentially relevant source of immunogenic cytoplasmic and nuclear self-antigens. Indeed, any abnormality in their clearance is potentially able to increase the availability of otherwise undetectable immunogenic auto-antigens through the induction of secondary necrosis or NETosis.

Annexin V/7-AAD profile examination of cells freshly isolated from the peritoneal cavity showed increased frequency of necrotic and apoptotic cells in pristane treated *Sparc*^*-/-*^ mice than WT counterpart (Figure 2A). This difference became significant after culturing PMN overnight (Figure 2A). Notably, the difference was restricted to the peritoneal cavity (the site of inflammation) and did not involve other organs, being neutrophil apoptosis and necrosis similar between Sparc deficient and proficient mice when evaluated in BM, spleen and PBL (Figure 2A). We then evaluated the capacity of *Sparc*^*-/-*^ and WT PMN to undergo NETosis. In agreement with our previous study performed in MRL-Lpr either crossed or not with *Sparc*^*-/-*^ mice (Sangaletti et al., 2014b), the PMN isolated from the peritoneal cavity of pristane-treated *Sparc*^*-/-*^ mice showed increased spontaneous capacity to extrude NET than similarly treated WT counterpart (Figure 2B).

**Figure 2:**
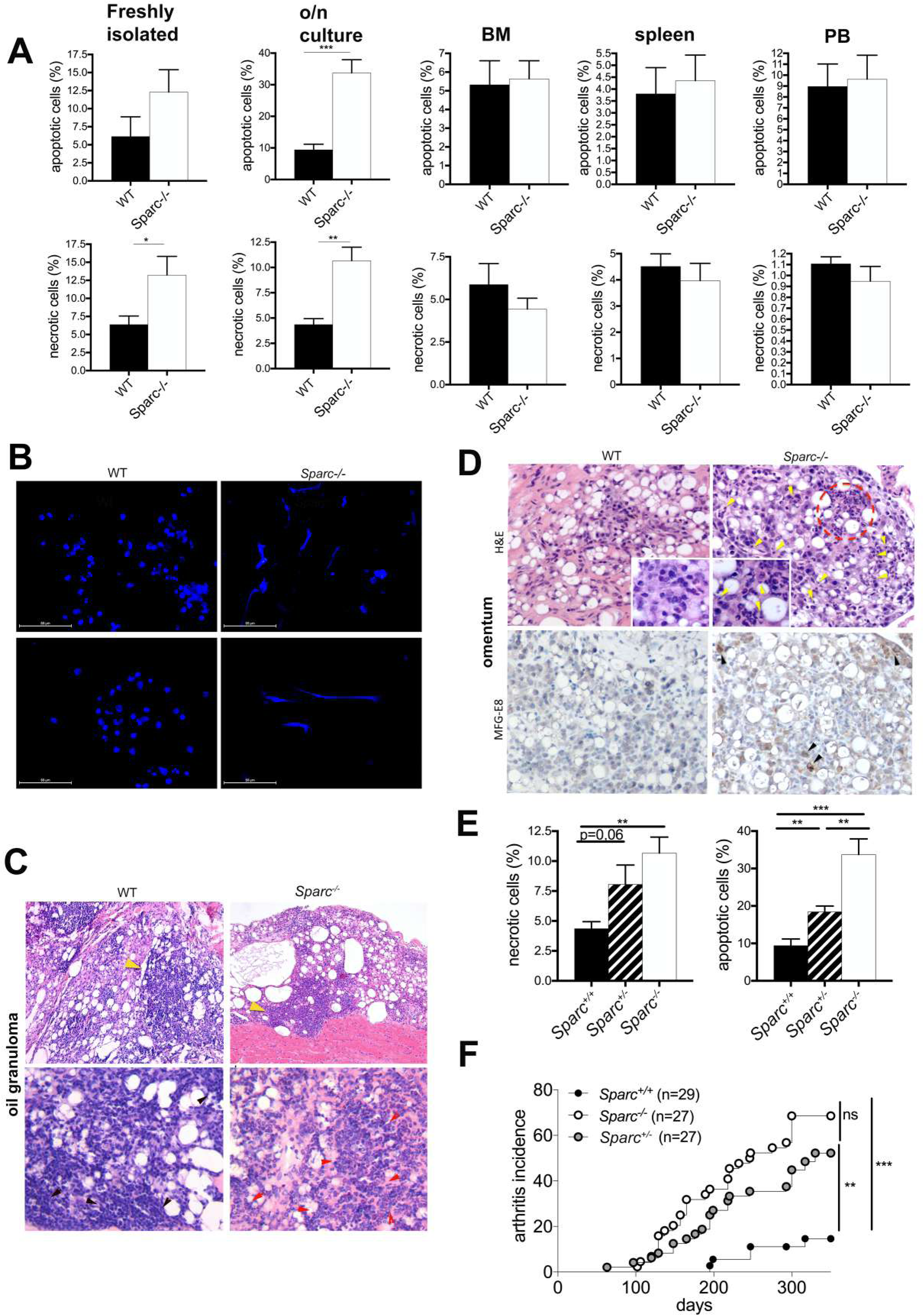
Accumulation of dying PMN in pristane-treated Sparc^-/-^ mice. A) Frequency of apoptosis (Annexin V^+^7AAD^+^) and necrosis (Annexin V^-^7AAD^+^) in PMN isolated from the peritoneal wash of pristane-treated WT and *Sparc*^*-/-*^ mice (day 130) showing a preferential accumulation of necrotic PMN and apoptotic PMN in *Sparc*^*-/-*^ mice; the same analysis, performed on the same cells cultured o.n., shows an amplification of the trend observed in freshly isolated PMN; the frequency of apoptotic and necrotic PMN isolated from the BM, spleen and PB from the same mice (n=7/group, representative of 5 experiments); B) NET formation by peritoneal PMN isolated from WT and *Sparc*^*-/-*^ mice mice (day 130 after pristane treatment). PMN were seeded onto poly-D-lysine coated glasses without any additional stimuli (Dapi staining); C) H&E staining of OG sections of the indicated mice at day 200 post pristane showing enrichment of PMN in OG from SPARC deficient mice; D) H&E staining (Upper panels) and MFG-E8 (Lower panels) of Omentum sections showing the predominance of lymphoid cells in WT mice conversely to the enrichment in PMN foci in *Sparc*^*-/-*^ mice (red dotted circle). Omentum from *Sparc*^*-/-*^ mice also shows a massive presence of cellular debris (yellow arrowheads) despite the presence in the same areas of Mϕ producing MFG-E8 (lower panel, back arrowheads); E) Frequency of apoptosis (Annexin V^+^7AAD^+^) and necrosis (Annexin V^-^7AAD^+^) in PMN isolated from the peritoneal wash of pristane-treated heterozygous *Sparc*^*+/-*^ mice compared to WT (*Sparc*^*+/+*^) and *Sparc*^*-/-*^ counterpart (day 130). F) Kinetic analysis of arthritis incidence in *Sparc*^*+/-*^ mice compared to WT (*Sparc*^*+/+*^) and *Sparc*^*-/-*^ mice (** p<0.01; *** p<0.001).

In addition, abundant auto-antigens can be found in tertiary ectopic lymphoid structures in form of oil granulomas (OGs) that develop in the peritoneal cavity of pristane-treated mice and that represent the primary sites of autoreactive B cells activation in this model. Also, OG (Figure 2C) and omentum (Figure 2D) were mainly populated by either lymphoid cells, or PMN in WT and *Sparc*^*-/-*^ mice, respectively. In the *Sparc*^*-/-*^ omentum we identified consistent numbers of cellular debris (Figure 2D, upper panels, arrowheads) apoptotic cells and the accumulation of macrophages competent for phagocytosis (MFG-E8 staining in Figure 2D, lower panels).

Overall these evidences suggest that the prevalent immunogenic cell death of *Sparc-/-* PMN and their specific enrichment in ectopic lymphoid structures are supportive for increased availability of auto-antigen and their presentation in *Sparc*^*-/-*^ mice. Corroborating this hypothesis, OG from pristane-treated *Sparc*^*-/-*^ mice showed a higher incidence of pseudo-follicular structures that are suggestive of follicular B cell responses.

Susceptibility to PMN death is a known hallmark of SLE, and since our data indicate that SPARC deficiency predispose to neutrophil death and SLE development, we analyzed peritoneal wash of pristane-treated hemizygous *Sparc*^*+/-*^ mice, which showed increased PMN death than WT counterpart (Figure 2E). Notably, hemizygous *Sparc*^*+/-*^ mice showed an intermediate disease score between WT and *Sparc*^*-/-*^ settings, in response to pristane treatment (Figure 2F).

### Reduced clearance of dying PMN in Sparc-/- mice is associated with defective modulation of “don’t-eat-me” signals

Considering that markers of PMN activation such as CD62L and ROS were equal in *Sparc*^*-/-*^ and WT PMN upon PMA stimulation (Figure 3A), we tested whether the increased accumulation of dying PMN in *Sparc*^*-/-*^ mice depend on their defective or incomplete removal by phagocytes. WT and *Sparc*^*-/-*^ thioglycollate-elicited macrophages (Mϕ) were cultured 20 minutes with PMN isolated ex-vivo from the peritoneal cavity of WT or *Sparc*^*-/-*^ mice, 130 days after pristane-treatment. To make a direct comparison, PMN were stained with distinct vital dyes (CFSE for WT PMN and Cell Proliferation Dye eFluor 670 for *Sparc*^*-/-*^ PMN) and mixed at 1:1 ratio before coculture with macrophages. PMN from pristane-treated *Sparc*^*-/-*^ mice were less efficiently phagocytosed than the WT counterparts (Figure 3B). To gain a rough information on the phagocytic activity of *Sparc* proficient and deficient macrophage, *in vivo* phagocytosis assay was performed by i.p. injection of UV-irradiated and CFSE labeled apoptotic Jurkat cells into thioglycollate-treated Sparc deficient or sufficient mice: results showed that both resident large peritoneal Mϕ and F4/80int monocyte-derived migratory Mϕ(Ghosn et al., 2010) similarly engulfed apoptotic cells, regardless the *Sparc* genotype (Figure 3D). Overall these data suggest that SPARC deficiency, per se, does not affect Mϕ efferocytosis but rather that the clearance of *Sparc*^*-/-*^ PMN is selectively defective. To confirm this possibility, we performed o/n time-lapse confocal microscopy analysis of PMN–macrophage co-cultures, documenting along time the interaction between macrophages and PMN as well as PMN cell death and their removal by Mϕ. Specifically, WT and *Sparc*^*-/-*^ PMN were labeled with the vital dye PKH26, and cultured with WT or *Sparc-*^*/-*^ macrophages in all possible combinations (Figure 3E). Differently from WT PMN that were almost completely engulfed by macrophages regardless the *Sparc* Mϕ genotype, *Sparc*^*-/-*^ PMN persisted in the co-culture.

**Figure 3:**
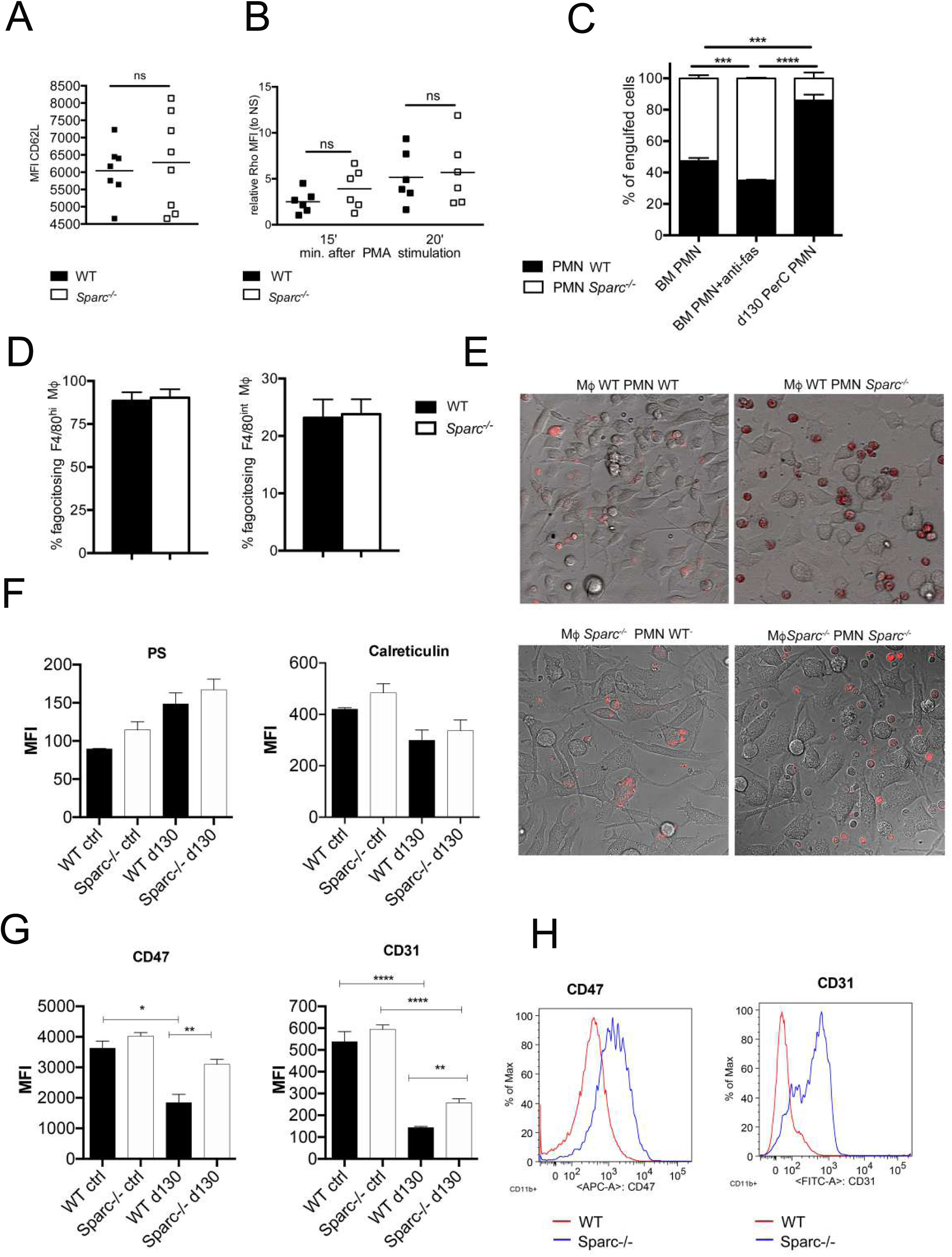
SPARC deficiency associates to a defective clearance by Macrophages of dying PMN. A) Cumulative FACS analysis showing CD62L surface level on freshly isolated peritoneal PMN and B) ROS production following *ex vivo* stimulation with PMA for the indicated times. All experiments were replicated at least 3 times (n=6/group). C) *In vitro* efferocytosis assay using WT peritoneal macrophages cultured 30 minutes with 1:1 ratio of WT and *Sparc*^*-/-*^ peritoneal PMN or BM PMN (treated or not with anti-Fas Ab, to induce apoptosis, as control). The analysis highlights a specific defect in the clearance of *Sparc*^*-/-*^ PerC PMN; D) % of F4/80hi and F4/80int peritoneal macrophages engulfing CFSE labeled apoptotic Jurkat cells; E) Time laps confocal microscopy of WT and *Sparc*^*-/-*^ macrophages (unstained, Bright field) co-cultured with either WT or *Sparc*^*-/-*^ macrophages (labeled with PKH26). Contrary to WT PMN that died and were almost completely engulfed by macrophages from both strains, *Sparc*^*-/-*^ PMN accumulated in the co-culture without removal; F) Expression of the “eat-me” (phosphatidylserine, PS, and calreticulin) and “don’t-eat-me” (CD47 and CD31) signals in peritoneal WT and *Sparc*^*-/-*^ PMN at day 130 after pristane treatment); E) Representative histogram plots showing the increased expression of CD47 and CD31 on *Sparc*^*-/-*^ PMN compared to the WT counterpart (pristane treated day 130).

Timelapse analysis clearly showed that the addition of *Sparc*^*-/-*^ PMN to WT macrophages was sufficient to block macrophage movement, impeding contact with PMN. On the contrary, macrophages move rapidly, chase and engulf WT PMN (Supplementary video 1&2). These results suggested a defect in *Sparc*^*-/-*^ PMN recognition. On this hypothesis, we evaluated the expression of “eat-me” (calreticulin and phosphatidylserine) and “don’t-eat-me” (CD47, CD31) signals (Bratton and Henson, 2011; Hart et al., 2000; Ravichandran, 2011) on PMN isolated from the peritoneal cavity of WT and *Sparc*^*-/-*^ pristane-treated mice (day 130). *Sparc*^*-/-*^ and WT PMN undergoing cell death showed similar expression of calreticulin and phosphatidylserine “eat me” signals (Figure 3F), however this expression in *Sparc*^*-/-*^ mice was not paralleled by a corresponding down-regulation of “don’t-eat-me” signals, particularly of CD47. Indeed, the expression of CD47 was retained and even up-regulated in *Sparc*^*-/-*^ PMN as compared to WT PMN (Figure 3G-H). Analyzing CD31, another don’t eat me signal, we found a similar trend with *Sparc-/-* PMN retaining CD31 expression higher than the WT counterpart (Figure 3G-H) This finding indicated a defective delivery of “eat me” and “don’t eat me” signals, in *Sparc*^*-/-*^ PMN, associated with their impaired clearance.

### *Dying PMNs that escape* Mϕ scavenging *are source of autoantigens for DC presentation*

Having shown that, in *Sparc*^*-/-*^ mice, the defective clearance of dying PMN provides a great source of auto-antigens, we explored whether *Sparc-/-*macrophages, although competent for the phagocytic activity, could contribute in worsening the autoimmune phenotype in *Sparc-/-* mice, supporting inflammatory pathways or APC functions. As first approach, we compared the gene expression profiles (GEP) of Mϕ isolated from the peritoneal cavity of pristane-treated WT and *Sparc*^*-/-*^ mice. Thirteen genes were up-regulated and 8 were down-regulated in *Sparc*^*-/-*^ macrophages (Figure 4A and Supplementary Table 1). Gene set enrichment analysis (GSEA) revealed up-regulation of gene sets proper of an inflammatory status, including TNF, IFNγ and IFNα (Figure 4B, Supplementary table 2). Furthermore, metabolic pathways relevant for macrophages activity, such as cholesterol homeostasis (relevant for M1) and oxidative phosphorylation (relevant for M2), were up or down-modulated, respectively, in *Sparc*^*-/-*^Mϕ. These data support a preferential polarization toward an M1 phenotype (Diskin C. 2018). In line, the GEP analysis shows that *Sparc*-deficient macrophages express significantly less Mrc1/CD206 than WT macrophages (FC= - 2,023; p<0.05, Supplementary Table 1)

**Figure 4:**
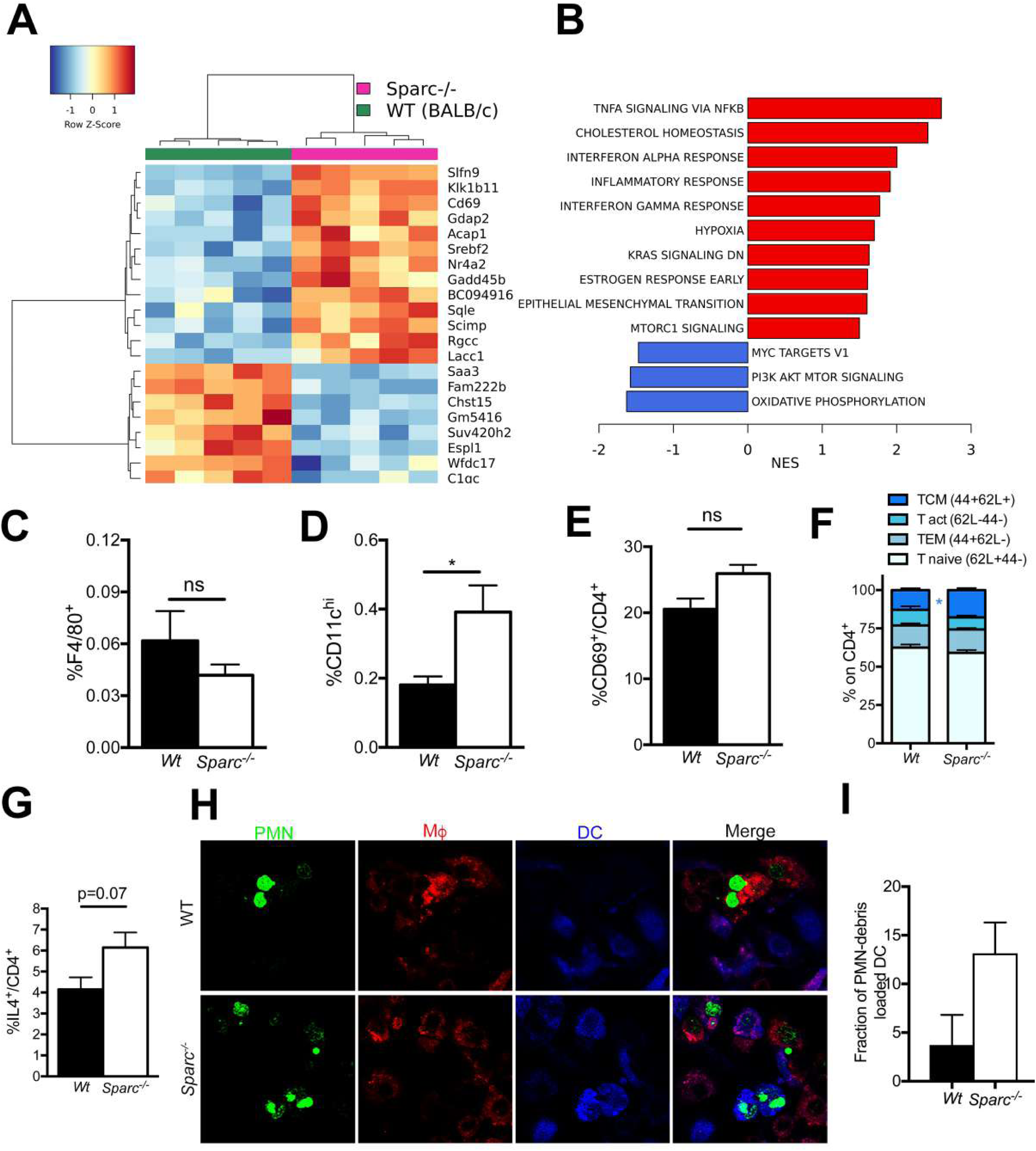
Increased DC activation and presentation in Sparc^-/-^ mice. A) Heat map of Mϕ isolated from the peritoneal cavity of pristane-treated WT in comparison to *Sparc*^*-/-*^ mice; B) Gene set enrichment analysis of WT and *Sparc*^*-/-*^Mϕ C) Frequency of F4/80+ Mϕ and D) of dendritic cells (CD11c^hi^) in mediastinal LN of pristane-treated WT and *Sparc*^*-/-*^ mice (* p< 0.05, Student *t*-test). E) Frequency of activated CD4^+^ T cells expressing CD69^+^ in draining mediastinal LN of pristane-treated WT and *Sparc*^*-/-*^ mice (** p< 0.01, Student *t*-test). F) Frequency of central memory (TCM CD44+CD62L+), activated (T act CD44-CD62L-), effector memory (TEM CD44+CD62L-) and naive CD4+ (CD44-CD62L+) cells in WT and *Sparc-/-* mice. G) Frequency of IL4+-CD4 T cells; H) Representative confocal microscopy analysis showing the preferential uptake of PMN debris by DC (blue) but not macrophages (red) in the *Sparc-/-* setting. I) Cumulative data obtained analyzing 5 sample/group for a total of n=50 DC and n=50 macrophages.

This M1 deflected pro-inflammatory microenvironment suggested to investigate the APC functions of macrophages and DC from *Sparc*^*-/-*^ mice. The frequency of F4/80+ Mϕ in draining LN was similar in both *Sparc* proficient and deficient mice (Figure 4C), however a significantly higher frequency of dendritic cells (DC, CD11c^hi^) was detected in *Sparc*^*-/-*^ LN (Figure 4D), supporting the involvement of DC in antigen presentation and T cells activation. In line, draining LN of pristane-treated *Sparc*^*-/-*^ animals showed an increased frequency of activated CD4^+^ T cells skewed toward a central memory T cell phenotype (Figure 4E-F). Although, not statistically significant, a trend towards increased IL-4 production by total CD4+ T cells was also observed (Figure 4G). This particular evidence finds correlation with SLE patients (Banchereau and Pascual, 2006) and is in line with the increased titer of T-cell dependent Abs to dsDNA (Rekvig and Nossent, 2003) observed in *Sparc*^*-/-*^ mice.

Overall, these results suggest that defective clearance of *Sparc*^*-/-*^ PMN by macrophages could increase the fraction of auto-antigens available for presentation by DC. To gain more insight into this point, we modeled the crosstalk of dying PMN with Mϕ and DC, *in vitro*. Overnight time lapse confocal microscopy was set up co-culturing combinations of bone marrow (BM)-derived Mϕ and BM-derived DC together with PMN isolated from the peritoneal cavity of pristane-treated mice. When PMN, Mϕ and DC were from WT mice, we observed that PMN progressively underwent apoptosis and were readily engulfed by Mϕ but not from DC. On the contrary, when the three populations were from *Sparc*^*-/-*^ mice, a relevant number of residual non-phagocytosed PMN was observed, and a notable fraction of PNM materials was uploaded into DC (Figure 4H-I).

This experiment underlined that the pro-inflammatory phenotype of *Sparc*^*-/-*^ Mϕ combined with a defective clearance of dying PMN could concur to the shift from a tolerogenic to an immunogenic environment licensing DC presentation.

### Accumulation and presentation of PMN-derived antigens contribute to the arthritic phenotype

Arthritis is the most relevant phenotype occurring in pristane-treated *Sparc*^*-/-*^ mice. Considering the prototypical proinflammatory cytokines IL17, IFNγ, and TNF involved in the pathogenesis of arthritis, we evaluated their production in pristane-treated WT and *Sparc*^*-/-*^ popliteal LN draining paws at day 130, a time point preceding overt tissue damage. We found increased IL17 and IFNγ in *Sparc*^*-/-*^ LN, whereas TNF although increasing, did not reach statistically significant difference with the WT counterpart (Figure 5A-B). In the inflamed joints, the expression of *Il17f* transcript was higher in *Sparc*^*-/-*^ than WT mice (Figure 5C). The source of IL17 was identified in T cells (CD4+), PMN (CD11b+Gr-1+) and γδ T cells (CD11b-CD4-γδTCR+) (Figure 5D), with the latter population contributing substantially. Indeed, although the frequency of γδ T cells was relatively low (between 0,5 to 3%), they were the most potent producers of IL-17 (Figure 5E). The increased granulocytic infiltration (Gr-1+ elements) observed histologically in *Sparc*^*-/-*^ paws, were consistent with an IL17-skewed inflammatory environment and suggested of evaluating whether PMN could represent a stimulus for γδ T cells either directly or through antigen presenting cells. To address this point, we set up co-cultures experiments in which CFSE-labeled γδ T cells were cultured with DC and PMN from WT or *Sparc*^*-/-*^ mice. We found that the concomitant presence of DC and PMN was required for effective stimulation of γδ T cell proliferation in WT conditions. However, when PMN were from *Sparc*^*-/-*^ mice they showed a direct capacity of stimulating γδ T cell proliferation, efficiently and independently of the presence of DC (Figure 5G).

**Figure 5:**
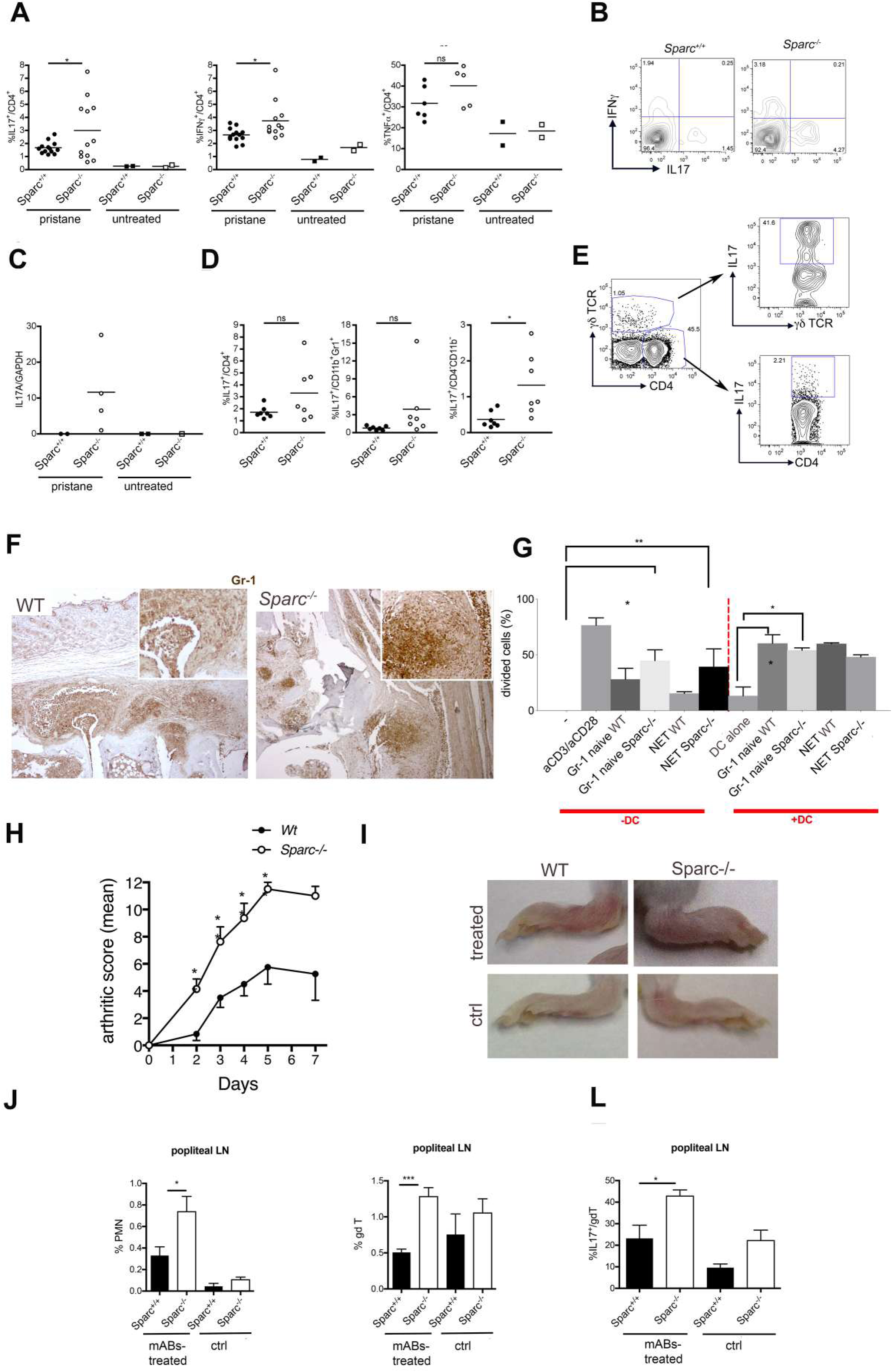
Increased IL-17 production in arthritic paws from Sparc^-/-^ mice. A) Intracellular staining for IL-17, IFN-γ and TNF-α in CD4^+^ T cells from draining LN of naive and pristane treated mice (130 days after treatment). Results are representative of three independent experiments; (* p< 0.05; Student *t*-test) B) Representative dot plots for the intracellular staining; C) qPCR for IL17A performing by digesting the whole paws of the indicated mice; results are normalized to GAPDH; representative of two other independent experiments; D) Intracellular staining for IL-17 in Th cells (CD45^+^CD4^+^), PMN (CD45^+^CD11b^+^Gr1^hi^) and other infiltrating cells (CD45^+^CD11b^-^CD4^-^) of paws isolated and digested from mice 130 days after pristane injection. E) Production of IL-17 by γδTCR+ and CD4 cells. Representative plots. F) IHC analysis for Gr-1 performed on paws obtained from WT and Sparc-/- mice. G) Frequency of proliferating CFSE-γδ T cells when co-cultured with DC and PMN either from WT or *Sparc*^*-/-*^ mice. This analysis highlights a direct capacity of *Sparc*^*-/-*^ PMN (either naïve or undergoing NETosis) of stimulating γδ T cells, independently from the presence of DC (* p< 0.05; ** p< 0.01, Student *t*-test). H) Arthritis score obtained evaluating anti-collagen II mABs-treated mice every two weeks (n= 17 mice/group pooled from two independent experiments); I) Representative pictures of paws from WT and *Sparc*^*-/-*^ at day 200 treated with anti-collagen II mABs. K) Frequency of PMN; L) γδ-T cells and M) IL-17+ γδ T-cells in mice treated with anti-collagen II mABs (* p< 0.05;** p< 0.01, ***p< 0.001; Student *t*-test).

These data would suggest that the more severe arthritis observed in *Sparc*^*-/-*^ than WT mice, other than result from accelerated onset of SLE-like autoimmunity, can be exacerbated by the local accumulation of PMN and their stimulation of IL17-producing γδ T cell. To further challenge this hypothesis, we induced arthritis in WT and *Sparc*^*-/-*^ mice adopting a mAb-induced arthritis model, a system that uses a pool of four anti-collagen II mAbs, bypassing, *de facto*, the contribution of systemic SLE-like autoimmunity to generate arthritis. In the mAb-induced model, *Sparc*^*-/-*^ mice developed arthritis more rapidly than their WT counterpart (Figure 5H-I) and showed popliteal LN enlargement (not shown) and PMN and γδ T-cell enrichment (Figure 5 J-K), with significant increased production of IL17 (Figure 5L).

Finally, as pristane-induced arthritis in mice surrogate human SLE-associated arthritis we tested the consistency of SPARC expression in human evaluating published gene expression profiles (GSE 36700) of synovial biopsies of knees affected by Rheumatoid Arthritis, Osteoarthritis, Seronegative Arthritis and SLE-associated Arthritis. ANOVA test was performed to compare each class of disease and SPARC had a p-value < 0.05 showing an overall change of expression with the lower expression in SLE patients who showed an opposite trend for EPSTI1 and MX1 as representative key genes in autoimmunity (Figure 6). This result underscores the possible contribution of SPARC down-regulation in the pathogenesis of SLE and arthritis.

**Figure 6:**
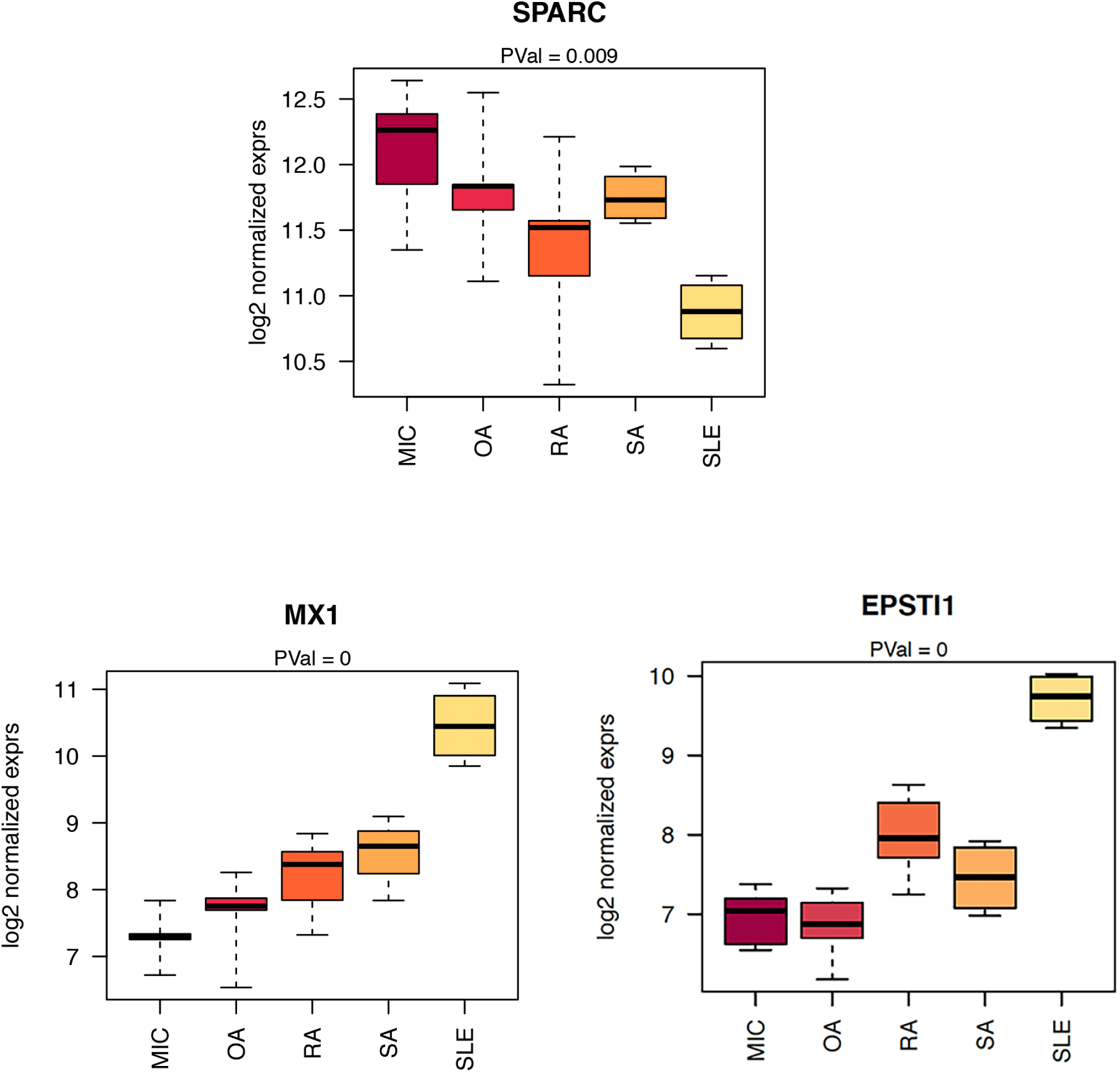
A) Expression of SPARC, MX1 and EPSTI1 in each class of disease compared with ANOVA-test (significance with p-value < 0.05);

## Discussion

This study unveils a new immunological function of SPARC controlling the correct removal of dying PMN, which otherwise provide a source of nuclear and cytoplasmic auto-antigens for the development of autoimmunity.

Using the pristane-induced SLE-like disease (Satoh and Reeves, 1994), we show that *Sparc*^*-/-*^ mice had anticipated and exacerbated SLE-associated autoimmune manifestations, despite reduced formation of ascites. This last observation was surprising since ascites has been considered a necessary step for generation of auto-antibodies upon pristane challenge (Satoh and Reeves, 1994). Nevertheless, this event seems unnecessary in *Sparc*^*-/-*^ mice, because of more efficient DC migration and antigen presentation and increased formation of tertiary lymphoid organs, in form of oil granuloma (OG), which are instrumental in the pathogenesis of systemic autoimmunity (Nacionales et al., 2006). In comparison to the *Sparc*-proficient counterpart, OGs from *Sparc*^*-/-*^ mice show a preferential accumulation of PMN associated with the presence of cell debris suggestive of active cell death. Accordingly, the peritoneal wash of pristane-treated *Sparc*^*-/-*^ mice showed increased numbers of dying PMN, which represent a candidate source of autoantigens. Although exposed to the same pristane treatment, *Sparc*-proficient PMN mostly die by apoptosis, whereas *Sparc*^*-/-*^ PMN die preferentially by necrosis and NETosis. It might be possible that SPARC deficiency could modify the outcome of cell death programs allowing PMN, not yet committed toward apoptosis, to release NETs. In this context, we have previously reported that complement deposition and activation (C1q and C5a) is increased in Sparc-deficient mice (Tripodo et al., 2017) and that C5a is both an inhibitor of apoptosis (Fishelson et al., 2001) and an inducer of NETosis in presence of IFNs (Yousefi et al., 2009).

Another interesting hint comes from the observed robust expression of “don’t eat me” signals: CD47 and CD31, in *Sparc*^-/-^ PMN, which could unbalance PMN death toward secondary necrosis and NETosis, in the pristane model. Also in cancer, CD47 overexpression has been associated with decreased PMN apoptosis and phagocytosis (Barrera et al., 2017).

The increased NETosis of *Sparc*^*-/-*^ mice could also account for the higher titer of a-dsDNA autoantibodies. Differently from a-RNP Ab, dsDNA Abs require IL-6 and develop according to the inflammatory peaks of disease ((Richards et al., 1998)). Although IL-6 is not directly involved in NET formation, the preferential and persistent interaction between NET and DC promotes DC activation and their release of IL-6, toward ds-DNA Ab production (Sangaletti et al., 2012). In *Sparc*^*-/-*^ mice, the increased NETosis along with the defective macrophages clearance of dying PMN might favor the development of this class of auto-antibodies. In support of the DC centrality in this process, we show that DC, but not macrophages, migrate to draining lymph nodes upon pristane treatment. In vitro, co-culture of PMN, DC and macrophages shows that the combination of defective macrophages clearance and of increased NETosis, associated to the *Sparc*^*-/-*^ genotype, allow a more efficient upload of DC with NET components. The clearance of NET and the associated MPO involves the macrophage scavenger receptor CD206/MRC1 (Shepherd and Hoidal, 1990) (Gordon and Pluddemann, 2018). GEP analysis shows that *Sparc*-deficient macrophages express significantly less Mrc1/CD206 than the proficient counterpart. This is partially explained by the preferential M1-polarization of *Sparc*-null macrophage being CD206 expression almost entirely restricted to M2 macrophages. Such polarization is also supported by the increased expression of genes related to cholesterol homeostasis, which characterize the M1 phenotype. Conversely, M2 macrophages are more dependent on oxidative phosphorylation (OXPHOS) over glycolysis for ATP supply (Oren et al., 1963). Overall, the *Spar* deficient condition favors DC uptake of NET debris and their presentation toward a-dsDNA Abs development.

Although we demonstrated that *Sparc*-deficiency in mice recapitulates key features of neutrophil defects characterizing SLE patients (Kaplan, 2011), humans are genetically *Sparc*-proficient. Nevertheless, the persistent presence of IFNs characterizing the initial phase of SLE might create the conditions promoting local SPARC down-modulation. Notably, we have previously shown *Sparc* down-modulation in mice in which peripheral autoimmune conditions was induced (Tripodo et al., 2017). Also, the bone marrow of these mice was characterized by local IFNγ production, downmodulation of Sparc and collagen genes in mesenchymal cells (Tripodo et al., 2017).

Notably, a variety of α-ds-DNA Ab from SLE patients target ubiquitary mesenchymal/ECM antigens including a 44-amino-acid fragment of Hevin, a protein belonging to the osteonectin/BM-40/SPARC family. Our link between defective *Sparc* expression and autoimmunity opens new prospects on investigating whether a-ds-DNA Ab targeting SPARC might be part of the pathogenic events leading to autoimmunity and interfering with PMN homeostasis, the latter inspired by the reported capacity of autoantibodies, from SLE sera, to impair PMN functions (Tsai et al., 2019).

## Author contributions

S.S. and M.P.C. designed the research; S.S., L.B. A.G., D.L., B.B., P.P., performed the experiments, C.T. performed histopathological analysis, M.M., L.DC, M.D. generated and analyzed in silico data. S.S. and M.P.C wrote the manuscript.

## Acknowledgments

This work was supported the Italian Ministry of Health (Young Researcher Grant number GR-2013-02355637). The authors also thank Mrs. E. Grande for administrative support.

